# Dispersal across habitat boundaries: uncovering the demographic fates of populations in unsuitable habitat

**DOI:** 10.1101/2021.03.22.436493

**Authors:** Megan C. Szojka, Rachel M. Germain

## Abstract

Patchy landscapes are characterized by abrupt transitions among distinct habitat types, forcing species to cross habitat boundaries in order to spread. Since seed dispersal is a probabilistic process, with a kernel that decays with distance, most individuals will fail to reach new, suitable habitat. Although failed dispersers are presumed dead in population models, their demographic fates may not be so simple. If transient survival is possible within unsuitable habitat, then through time, individuals may be able to reach distant, suitable habitat, forming new populations and buffering species from extinction. In a fragmented Californian grassland, we explored the fates of individuals that crossed habitat boundaries, and if those fates differed among specialists dispersing from two habitat types: serpentine habitat patches and the invaded non-serpentine matrix. We surveyed the diversity of seedbank and adult life stages along transects that crossed boundaries between patches and the matrix. First, we considered how patch specialists might transiently survive in the matrix via seed dormancy or stepping-stone populations. Second, we investigated the dispersal of an invasive matrix specialist (*Avena fatua*) into patches, to assess if sink populations existed across the habitat boundary. We found that dormancy maintained populations of patch specialists deep into the matrix, as abundances of seedbanks and of adult plant communities differed with distance into the matrix. We found evidence that these dormant seeds disperse secondarily with vectors of material flows in the landscape, suggesting that they could eventually reach suitable patches even if they first land in the matrix. We found that *A. fatua* were largely absent deep in patches, where reproductive outputs plummeted and there was no evidence of a dormant seedbank. Our results not only reveal the demographic fates of individuals that land in unsuitable habitat, but that their ecological consequences differ depending on the direction by which the boundary is crossed (patch → matrix ≠ matrix → patch). Dormancy is often understood as a mechanism for persisting in face of temporal variability, but it may serve as a means of traversing unsuitable habitat in patchy systems, warranting its consideration in estimates of habitat connectivity.

## Introduction

Dispersal plays a central role in biodiversity maintenance across spatial scales, given that dispersal limitation is one of the main challenges populations face in patchy environments (Loreau et al. 2003, Carrara et al. 2012, Gilbert & Levine 2013, Germain et al. 2017, Lancaster & Downes 2017). In many ecological models, dispersal is treated as a rate of individuals leaving one habitat patch and entering another (i.e., island biogeography [MacArthur & Wilson 1963], metacommunity [Leibold et al. 2004], and metapopulation theories [Hanski 1982, Harrison 1991]). However, doing so assumes that individuals either never fail to disperse to suitable habitat, that failed dispersal results in immediate death, or that only dispersal that has been successfully realized is important (Fahrig & Merriam 1985, Pulliam 2000). Modeling dispersal as a realized rate is reasonable for many organisms, such as species with active dispersal who easily traverse a mosaic of habitat types (e.g., boreal birds [Gobeil and Villard 2002], butterfly distributions [Dapporto 2010]), yet for many others, individuals frequently disperse into unsuitable habitat (Cook et al. 2002). If an individual lands in unsuitable habitat, three demographic fates are possible: the individual either dies, transiently survives in a dormant state, or transiently survival in a non-dormant state with some low probability of reproducing at all. For those species that are forced to disperse across habitat boundaries, transient survival may be particularly critical for species’ long-term persistence in a landscape (Leibold et al. 2004, Chisholm et al. 2011, Snyder 2011, Gilbert & Levine 2013, Wisnoski et al. 2020).

As individuals disperse from patches into the matrix, remaining dormant is one demographic fate which might increase future opportunities to disperse to new patches (Wisnoski et al. 2019). To understand why, we first discuss the most probable fate of an individual dispersing in a fragmented system. Dispersal is a process governed by chance, where a probability density function, or ‘dispersal kernel’, describes the decaying probability of landing at increasing distances away from a parent plant (Nathan 2006). As a result, although an individual which leaves a patch has some rare chance of reaching a new, far away patch (Snyder & Chesson 2003), it is far more likely that the individual will land short, in the unsuitable habitat (Ismail et al. 2017). If that individual is unable to become dormant, it would fail to survive or to reproduce and would therefore fail to contribute to population dynamics in the future (Harrison 1991). Although dormancy and dispersal are often considered distinct strategies for coping with different axes of environmental variation (i.e., temporal vs. spatial, respectively; Snyder 2006), dormant individuals may also disperse. Indeed, secondary dispersal is common in nature, for example, plant seeds which require passage through an animal’s gut to break dormancy, meaning that dormancy and dispersal are necessarily coupled at the individual level, and can affect species distributions (Hämäläinen et al. 2017, Penfield 2017). If dormant individuals do disperse, individuals may be able to permeate the unsuitable matrix, increasing opportunities to access habitat patches at distances beyond their own intrinsic dispersal abilities in ways not currently considered by metapopulation models (Padilla et al. 2012, Snell Taylor et al. 2018).

Species which specialize on matrix habitat also disperse, and if they land in unsuitable patches, matrix specialists could have demographic fates and ecological consequences that are different compared to those of patch specialists that enter the matrix. Here we consider two differences. First, rates of dispersal from matrix habitat into habitat patches tend to be higher than the reverse. Matrix habitat is continuous by definition and often covers a large area relative to habitat patches (Cook et al. 2002), supporting high population abundances. Second, in many systems, such as those characterized by invasion, matrix habitat forms in more productive portions of the landscape, resulting in high *per capita* seed outputs (i.e., for plants; Rand et al. 2006). Together, high abundances and high *per capita* seed outputs mean that matrix specialists frequently disperse into non-matrix habitat, forming edge effects (Harrison et al. 2001). If high numbers of dispersing seeds germinate into adults, those adults could competitively impact diversity in patches even if they only exist as transient sink populations (λ<1) with reduced *per capita* competitive effects (Hart et al. 2018). Alternatively, dispersing seeds may never germinate, in which case spillover of matrix specialists would minimally impact species in patches even if they accumulate in the dormant seedbank (Wisnoski et al. 2019). Although past studies show that matrix species can occur in patches at high abundances due to seed spillover (Harrison et al. 2001), none to our knowledge have tracked the demographic fates of those individuals or explicitly compared rates of spillover in both directions (matrix → patch vs. patch → matrix). Doing so is worthwhile as these differences translate into different consequences for persistence of patch specialists, matrix specialists, and the overall composition of biodiversity as species cross habitat boundaries.

Not all habitat boundaries are created equal. Rates of dispersal depend on how species’ dispersal traits interact with landscape features (Germain et al. 2019). Specifically, many species possess specific combinations of traits, known as ‘dispersal syndromes’ (Howe & Smallwood 1982), that are specialized to facilitate dispersal via particular dispersal vectors, such as gravity (e.g., spherical seeds move downhill farther than flat seeds) or animals (e.g., burrs or awns that facilitate attachment; Römermann et al. 2005, Padilla et al. 2012). Yet dispersal vectors do not intersect all patches equally—rather, they vary spatially due to the physical structure of landscapes, which directs material flows or behavioural decisions made by animals (Dunning et al. 1992, Clobert et al. 2009). Consequently, we expect that dispersal kernels (and thus a population’s ability to permeate the matrix) will also vary spatially, stretched outward for species with dispersal syndromes in locations strongly intersected by their corresponding dispersal vectors. This contrasts how dispersal is treated in ecological research - as a stochastic external process unaffected by species’ biologies or landscape context (but see Lowe & McPeek 2014).

In a serpentine grassland, our aim is to identify whether populations are transiently maintained in unsuitable habitats, and if so, which demographic stage facilitates this survival. This study is the first step towards determining whether transient dispersers can contribute to regional dynamics by eventually ending up in suitable habitat. Serpentine grassland communities are an ideal system to test spatial biodiversity questions as they occur as discrete patches embedded within an invaded habitat matrix. We hypothesized that (i) patch specialists that disperse into the invaded matrix persist as dormant seeds, (ii) habitat boundaries with dispersal vectors that correspond with a given dispersal syndrome may facilitate the dispersal of dormant seeds and (iii) spillover and germination of an invasive matrix specialist maintains sink populations across habitat boundaries. We sampled plant communities across the habitat boundaries between patches and the matrix, comparing diversity between the seedbank (all viable seed present prior to the growing season, including those that will remain dormant during the growing season) and the germinated adult community (emerging from the non-dormant fraction of the seedbank). If seed dormancy allows patch specialists to persist within the matrix (hyp. i), then we predict that the decline in species richness and total abundance at increasing distances into the matrix will be more pronounced for the adult community than for the seedbank. If dispersal vectors stretch the dispersal distances of dormant seeds farther into the matrix (hyp. ii), then we predict that decline in total seed abundance with increasing distance into the matrix will be weakest where habitat boundaries are traversed by specific dispersal vectors (e.g., animal-dispersed species along animal paths), compared to habitat boundaries where dispersal vectors are either absent or do not align with species’ dispersal modes (e.g., wind-dispersed species along animal paths). Lastly, if matrix specialists form sink populations in habitat patches (hyp. iii), then we expect to find many individuals present as adults and few as seeds, with adult reproductive outputs below replacement (i.e., mean per capita reproduction <1). As we will show, although patch specialists and matrix specialists both occur across habitat boundaries, their occurrences are driven by different mechanisms and differ in their implications for biodiversity maintenance.

## Materials and methods

### Study system and site selection

McLaughlin Natural Reserve in Northern California (38.860643 N, −122.364156 W) has a mediterranean-type climate, with cool wet winters (average rainfall of ∼750 mm, Germain et al. 2017) and hot dry summers. Within the reserve, we worked in serpentine grassland habitat, where the majority of plants have an annual life cycle (CalFlora 2019). Annual plants germinate with the initiation of the wet season (mid-October), followed by maturation, seed set, dispersal, then adult death through the summer (April − September).

Serpentine soils are geological outcroppings of the Earth’s mantle that are found along subduction zones (Brady et al. 2005). For that reason, they occur in a patchy configuration and are characterized by unique plant communities that are adapted to tolerate low nutrients and high heavy metal content (Kruckeberg 1954, Oze et al. 2008). The patches also act as a refuge from competitive exclusion by invasive plant species, notably, European grasses (e.g., *Avena fatua*), that cannot tolerate serpentine soils, and therefore dominate the surrounding non-serpentine matrix (Anacker 2014). As a consequence, habitat boundaries between patches and the matrix are characterized by abrupt environmental gradients, suggesting that dispersal into unsuitable habitat is common.

We haphazardly selected 11 serpentine patches out of 21 possible patches found within a 2.25 ha region of the reserve. All patches occurred on a south facing slope of a serpentine ridge, 469-530 m above sea level. We did not sample patches if they lacked abrupt habitat boundaries, as this feature was key to testing hypotheses about dispersal over a habitat edge. In this same region, past research has demonstrated that serpentine plant communities are dispersal limited (Germain et al. 2017) and that the degree of dispersal limitation is determined by landscape features (e.g., patch steepness, tree cover) which affects spatial patterns of wind, water, and animal movement (i.e., dispersal vectors; Germain et al. 2019).

To test how and why diversity changes across habitat boundaries, for each of the 11 patches, we sampled two to four transects that extended from within patches into the matrix. Each transect was oriented in a different direction on the hill face to systematically sample different possible dispersal pathways (lateral, uphill, or downhill, and following animal paths). Patches differed in size, so in small patches (<10 m diameter), to avoid overlapping transects that lacked independence, we sampled only two transects. In large patches (>30 m diameter), overlapping transects were not a concern, so we sampled up to four transects. Each transect contained four 0.03 m x 0.3 m plots: one plot 1 m into the patch, one on the habitat boundary edge, one 1 m into the matrix, and one 5 m into the matrix. In addition to the four plots along each transect, a single 0.09 m^2^ plot was sampled in the center of each patch. Note that, since patch sizes varied, the distance of the center plot from the patch edge also varied. We recorded the GPS location of each plot. In total, there were 9-17 plots per patch (depending on the number of transects), totalling to 129 plots across all 11 patches.

### Survey of seedbank diversity

Our first goal was to examine how seed dormancy could allow for the persistence of patch specialists across the habitat boundary (hyp. i), thus prior to the growing season, we collected the seedbank in each plot to compare against diversity in plots that arose from seeds that germinated during the growing season (described in *Survey of adult plant diversity*). The seedbank includes all viable seed in a plot, including seed destined to be dormant and non-dormant once the growing season initiates. Because most plant species in our grassland study system have an annual life cycle, the entire community can be captured through the seedbank in the dry season. In November 2018, before the fall rains initiated the growing season, we collected the entire seedbank in each of the 129 plots, including all seeds that had been produced the previous year and all dormant seeds that were stored from previous years (as some seeds survive for multiple years). All soil material, including seeds and dry vegetation, were collected with a trowel and brush and were then stored in paper bags. Because seeds are too small and difficult to separate from non-seed material to identify by eye, seedbank samples were shipped to the University of British Columbia (Vancouver, Canada) to be identification.

We counted and identified germinants that emerged from the seedbank in order to test our predictions that the total abundance of seeds (hyp. i & iii), species richness (hyp. i), and the identity of species (hyp. ii) would vary across transects. In January 2019, we germinated a subset of each seedbank sample in a growth chamber. The chamber conditions were set to emulate a typical Mediterranean winter, maintaining a temperature of 12°C and a 12-hr light cycle. Twenty grams of each seedbank sample were added to round growing trays of Sunshine mix #3. Each tray was hand watered every other day as seedlings started to germinate, up until 15 days at which point watering was withheld for one week to coax remaining viable seeds to germinate.

Once new seedlings began to mature, we watered every four days to keep the soil damp and increased the temperature to 15°C. Once all plants were old enough to identify (after ∼one month), we clipped all standing biomass with garden shears and then allowed the soil to dry completely. We then treated the same soil samples with 500 ppm gibberellic acid (GA3, Sigma-Aldrich) to chemically break the dormancy of any ungerminated seeds and track their growth to maturity. Throughout the trials, we counted the number of individuals of each species in every tray every two weeks and took photos of any unknown individuals for later identification. Nearly all individuals were identified to the species level except for *Hesperolinon* sp., *Lactuca* sp., and *Hordeum* sp., which could only be identified to the genus level. For a complete species list see Appendix S1: Table S1.

### Survey of adult plant diversity

To compare the spatial distribution of plant diversity among adult plant communities and the seedbank (hyp. i-iii), we returned to McLaughlin Natural Reserve in May 2019 to survey plants that germinated naturally and survived to adulthood in the field during the growing season. These adult communities had therefore germinated naturally from the same generation as the seedbank community that we had collected. Because we had removed the entire seedbank from each plot the previous fall, rendering them devoid of plant matter, we positioned our survey plots immediately adjacent to the seedbank plots to record the percent cover of each adult plant species present within an equivalent area (i.e., 0.03 m x 0.3 m). This served to approximate the community of adults that would have germinated from a nearly identical seedbank. Throughout the manuscript, ‘seedbank’ refers to all seeds that we collected in the field, whereas ‘dormant seeds’ refers to mismatches between diversity in the seedbank and the adult community (i.e., no dormancy if they align perfectly, dormancy if they do not align)—note that, by alignment, we do not mean absolute numbers of individuals, but rather, how each responded to position along transects (described further in *Analyses*). Past work has demonstrated that many species which specialize on serpentine soils can form persistent populations in non-serpentine soils (here, the habitat matrix) in the absence of competitors, but often fail to do so as they are excluded by invasive matrix specialists (Safford et al. 2005, Gilbert & Levine 2013). Hypothesis (i) was concerned with comparing the germinated adult community against the abundance and richness of seedbank, so we did not collect fitness data to discern transient (seed production <1) from persistent (seed production >1) adult populations as this is a well-established feature of our serpentine grassland system.

We tested the spillover and persistence of a matrix specialist across habitat boundaries (hyp. iii) using *Avena fatua* (hereafter *‘Avena’* for brevity) as a representative species. *Avena* was proportionally the most abundant species in the matrix, which, relative to habitat patches, is a species-poor habitat (matrix specialists comprised 24% of the regional species pool). *Avena* made up 18.2% of the abundance of all matrix-dwelling individuals. To understand *Avena*’s potential impacts on patch diversity, we returned to McLaughlin Reserve in October 2019 to quantify *Avena*’s seed output (a component of individual fitness of annual plants). To do so, we revisited our field survey plots at two of the most sparsely vegetated study patches, representing the most resilient refuges to invasion. If these patches are invaded by *Avena*, we would expect that more productive patches would also be invaded. We note that through estimating *Avena* abundances through germinating seedbank samples we cannot estimate the abundance of inviable seeds and therefore do not attempt to disentangle dispersal limitation from seed death across the habitat boundary. At each of these two patches, we revisited the same transects described earlier (*n* = 4 transects, 17 plots per patch), counting the number of seeds that three randomly-selected *Avena* individuals produced. If no *Avena* individuals were present in a plot, we found the closest adult *Avena* that was the same distance from the edge, within 1 m of our original plots. Together across our surveys, we have estimates of how many *Avena* seeds lurked in the seedbank, how many were present as adults, and what seed outputs those adult individuals had on average—in other words, we can disentangle the demographic fates of individuals along transects.

### Analyses

Because our hypotheses concern species spillover between patch and matrix habitat, we categorized species based on habitat affinity: patch specialists or matrix specialists. To categorize species in this way, we leveraged a 17-year historical dataset of species occupancy in serpentine sites and in non-serpentine sites at McLaughlin Reserve (80 sites total, courtesy of S. Harrison; Appendix A). From these data, we created rank-abundance curves of species abundance in each habitat type (serpentine vs. non-serpentine) and used these curves to assign each of our species a level of patch specialization. We additionally performed a sensitivity analysis which showed that our results were qualitatively equivalent using different thresholds of habitat affinity (where more conservative thresholds contained fewer rare species, see Appendix A and Appendix S1: Table S2). We used species within sensitivity levels one and two in our analyses, excluding the rarest species.

For all analyses described below, we used the same general statistical approach and tools. Specifically, all analyses were done in R version 3.6.1 (R core team 2019) using generalized linear mixed effects (glme) models with package ‘glmmTMB’ (package; Brooks et al. 2017). Statistical significance of fixed factors in our glme models was tested using type III analysis of variance with the ‘car’ package, where the reported *P*-values throughout are calculated from chi-squared tests of maximum-likelihood ratios (Fox & Weisberg 2019). Additionally, for every model we will describe, we used a forward model selection procedure to compete models of increasing complexity (i.e., more parameters). Model fit was assessed using a corrected Akaike Information Criterion (AICc, ‘MuMIn’ package; Barton 2019), retaining models with increased complexity if ΔAICc was at least −2 (see Appendix S2: Table S1 for model selection procedure). Although forward stepwise model selection has been shown to inflate Type 1 errors (Mundry & Nunn 2009), we additionally compared the fits of our final models to the full models using ‘Anova’ to confirm that the final models provided a significantly better fit (Appendix S2: Table S5).

### Analyses to test hyp. i: demographic fates of patch specialists in the matrix

Before analyzing our data, we transformed some of our variables in order to aid comparisons among data types and ease model fitting. First, for feasibility reasons of counting individuals in the field, the total abundance of individuals in the seedbank and adult community were sampled in different units (number of individuals vs. percent cover, respectively). Additionally, we subsampled the seedbank community when setting up our germination trials (described above). As such, we transformed both variables to make them comparable, specifically by rescaling each to vary from 0 to 1. In doing so, differences in intercepts between variables are not biologically meaningful (e.g., driven by sampling effects), but our hypotheses concern differences among variables in how strongly they vary along transects (i.e., a difference in slope), which are comparable. Second, we exponentiated the variable ‘distance’ (i.e., position along a transect relative to the habitat boundary), after adding a constant so that all values were positive, to draw low values up towards the central distribution (the inverse of a log transformation); this simply introduces non-linearity to our relationships via a linear transformation. Note that the exponentiation of ‘distance’ was only necessary for hypothesis (i) analyses as the distances introduced by patch centre positions were ‘low’ values only when considering patch to matrix dispersal.

To test hypothesis (i), we performed two analyses, each with a generalized linear mixed effects (glme) model (‘glmmTMB’ package; Brooks et al. 2017, and ‘lme4’ package; Bates et al. 2015), with different response variables: species richness and total abundance of individuals. For both analyses, we started with the same base model: ‘response’ ∼ life stage*poly(exp(distance), 2) + (1|patch id) + (1|transect id). As such, each base model included the interaction between distance over habitat boundaries and life-stage (germination trial and in-field surveys) as it was central to our biological question of whether the seedbank differed from the adults in different positions for richness and total abundance. We used the ‘poly(distance, 2)’ function to improve fit (based on AICc model comparison). The ‘poly’ function differs from squaring distance (a raw polynomial) as it orthogonalizes first-order and second-order polynomials before regressing them, comparatively improving our computational inferences by avoiding collinearity (Kennedy & Gentle 1980). Due to their different data types, we used Poisson and Beta error distributions for species richness and total abundance, respectively. For both richness and total abundance, we tested the interaction between life stage (i.e., seedbanks or adults) and distance from the habitat edge along transects (a continuous variable, negative values when in patches and positive values in the matrix). Treating ‘distance’ as a continuous variable has the benefit of retaining relative distances of plots along the transects from the habitat boundary, but, may be biased if significant interactions were solely driven by differences within patches rather than differences in the matrix (as the latter was most relevant to our hypotheses). To account for this possibility, we re-ran our analyses with distance from the habitat boundary as a categorical variable with 5 levels (i.e., patch centre, 5m into patch, 1m into patch, edge, 1m into matrix, 5m into matrix). Our results were qualitatively similar for both versions of analyses (Appendix S2: Table S2), but we present our results with ‘distance’ as a continuous variable as differences among relationships are more intuitively interpretable. Transect and patch identity were included as random effects to account for our nested sampling design.

### Analyses to test hyp. ii: stretching of dispersal kernels by dispersal vectors

Plant species’ dispersal syndromes may affect the distance seeds are found from the patch edge due to their interaction with landscape features. We searched the functional trait literature to determine the primary dispersal mode of each species in our dataset, specifically using past work with some of the same species as a starting point (Jones et al. 2015) and through datasets compiled by the Centre for Agriculture and Bioscience International (Appendix S1: Table S1; CABI.org). We categorized patch specialists into three dispersal syndromes: self-dispersing dispersal (including all autochory strategies, such as ballistic and gravity dispersed seeds), wind-dispersed (seeds often characterized by small size and a pappus) and animal-dispersed seeds (including endozoochory and epizoochory). In total, at our study site, dispersal by animals was the most common dispersal syndrome exhibited by patch specialists (*n* = 33 species), followed by self-dispersing (*n* = 28), and lastly, dispersal by wind (*n* = 14).

After assigning each species a dispersal syndrome, we analysed whether changes in total abundance of seeds in the seedbank across habitat boundaries could be explained by dispersal syndromes and dispersal vectors (hyp. ii). To test this, we started with the following base model: seed abundance ∼poly(distance, 2)+(1|patch.id)+(1|transect.id), using a Poisson error distribution. To our base model, we used forward selection to test whether the following fixed factors improved model fit (see Appendix S2: Table S3 for procedure): dispersal syndrome, transect type and steepness. ‘Transect type’ describes how transects are positioned within patches, both in terms of likely material flows by gravity (i.e., uphill, downhill, lateral) and whether transects fall along animal paths, as these variables influence seed dispersal that may align with different dispersal syndromes (hyp. ii). ‘Steepness’ (measured at the scale of an entire patch) was included because it may influence the magnitude of change in abundance along these transect types (i.e., down or up a steep vs. shallow slope). Because we detected a significant interaction between dispersal syndrome, landscape steepness, and poly(distance, 2), we used post-hoc Tukey tests to compare differences in relationships among dispersal syndromes (‘emmeans’ package; Lenth 2019).

### Analyses to test hyp. iii: demographic fates of a matrix specialist in patches

Next, we tested how abundant *Avena* was in habitat patches as seeds and as adults across habitat boundaries, quantifying per capita seed production of adults to assess whether those adult individuals (if they occurred at all) were likely contributing to source or sink populations (hyp. iii). Because we germinated a subset of material collected from each plot, in order to estimate the total number of seeds each plot contained, we back-calculated the amount of seeds that germinated in our 20 g subsamples to the total weight of all material collected from a plot.

We built glme models, either with abundance or seed production as response variables, with the following base model: abundance∼life-stage*poly(distance, 2)+(1|patch.id)+ (1|transect.id) and seed.production∼poly(distance, 2)+(1|patch.id)+(1|transect.id). For model selection, we tested four possible fixed effects: patch area (m^2^), productivity (NDVI), transect type, and steepness. The ability of invasive species to persist in serpentine soils may be limited by the harshness of the soil, which is why we included ‘productivity’ as a fixed effect. To estimate productivity, we obtained each patch’s NDVI (normalized difference vegetative index), as well as patch area (m^2^) and steepness using GIS (USGS 2020). Our glme models with abundance and seed production were analyzed with negative binomial and Poisson error distributions, respectively, as these distributions provided the best fit to the data (based on AICc model comparison).

## Results

We identified 77 plant species spanning 22 families in our community surveys (*n*_*plot*_ = 129, *n*_*transect*_= 30, *n*_*patch*_ = 11), which is the regional pool of species observed in our study. Seventy-four percent of species from the regional species pool were represented in the seedbank, whereas 92% of the species pool were observed in the adult community; six species were found only in the seedbank and 19 species were only found as adults. Across habitat types, 44 species were associated with serpentine patch communities (‘patch specialists’) compared to 18 species that were associated with the matrix (‘matrix specialists’). Thirteen species were associated with neither habitat (‘unassociated species’), as they were either habitat generalists or too rare to be attributed to any particular habitat. The remaining two species could not be identified and were listed as unknown (Appendix S1: Table S1). All summary statistics are presented as mean ± SE. Below, we begin by first reporting our findings for patch specialists dispersing into the matrix, and then for the dispersal of *Avena* (an invasive matrix specialist) into patches.

### Failed dispersal into the matrix results in a dormant seedbank (hyp. i)

We found partial support of the hypothesis that, due to seed dormancy in unsuitable habitat, species richness and total abundance of patch specialists would decline into the matrix more strongly for the adults than the seedbank. Specifically, we found that patch specialists maintained higher total abundances in the seedbank compared to adults at increasing distances away from patches (significant interaction between poly(exp(distance), 2) and life stage (*χ2* = 6.75, *P* = 0.034; Fig. 1B)), but there was no evidence that declines in species richness across distance differed between life stages (non-significant interaction between poly(exp(distance), 2) and life stage (*χ2* = 1.35, *P* = 0.421; Fig. 1A)).

**Figure 1.**
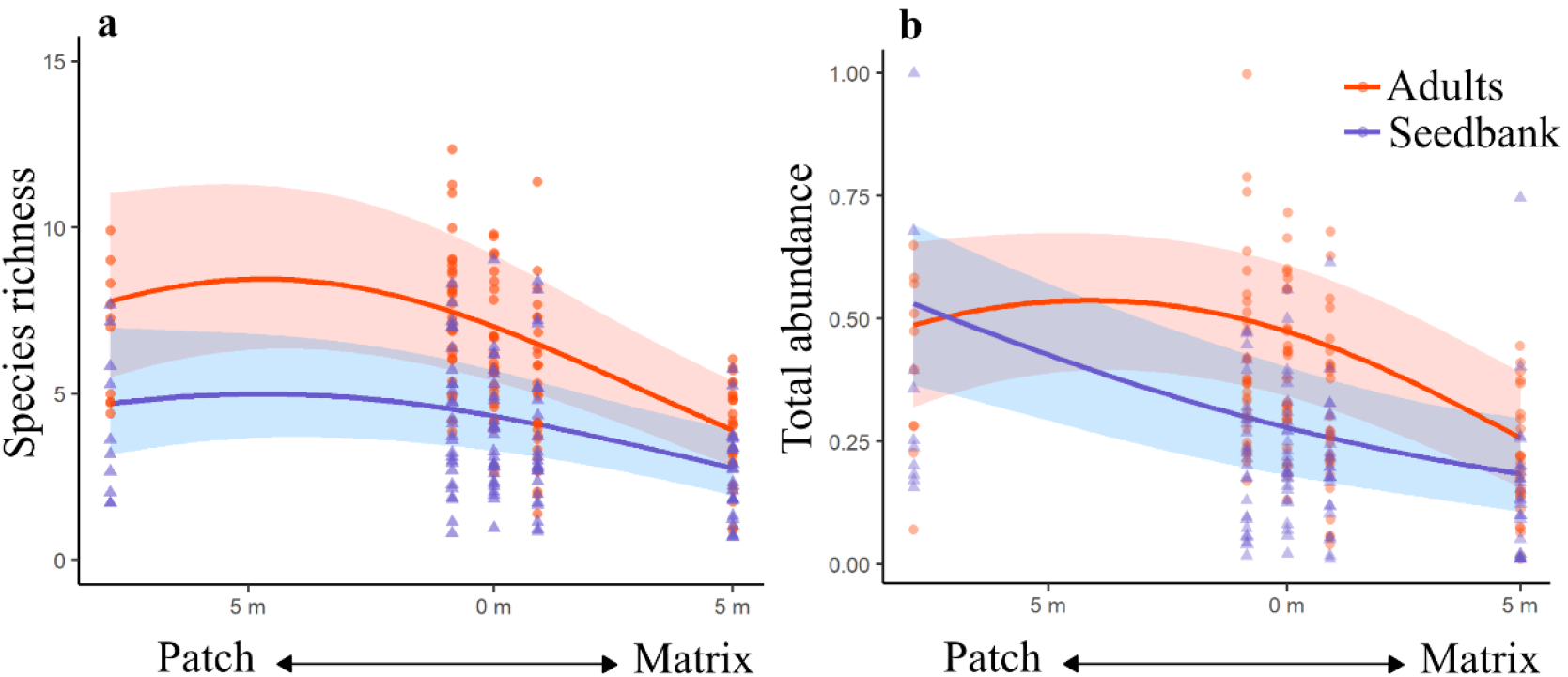
Effect of patch specialists crossing habitat boundaries on the structure of a) species richness and b) total abundance of adult vs. seedbank communities. The seedbank overall has lower richness and total abundance than the adult community; however, we were interested in how the slope of each metric differs between life-stages as they cross into the matrix. a) The change in richness with distance does not differ significantly between life stages. b) Total abundance is the sum of species abundances within each plot, which was then scaled for both the seedbank and the adults from 0 to 1 to compare the proportion of total abundance that is lost from the patch to the matrix for each life-stage. The change in total abundance differs between life stages: for every metre increase (from patch → matrix), the seedbank retains 4.02 ± 1.53, more individuals than the adult community (*P* = 0.008). Points are the raw data (*n* = 129), lines are fitted relationships from the models, and shaded areas are the 95% confidence bands of the model predictions. Points 8 m into the patch are plots at the patch centers, and since patches were different sizes, these range from 8-20 m from the boundaries.

### Landscape steepness, not dispersal vector, enhanced seed dispersal regardless of dispersal syndrome (hyp. ii)

We found support for our hypothesis that landscape features affect dispersal of patch specialists into the matrix, but not in the specific ways we expected. Specifically, contrary to our prediction that seed accumulation in the matrix would be highest in areas of the landscape where dispersal vectors aligned with dispersal syndromes, transect type (a proxy for dispersal vector) was not retained in our final model. Instead, only one landscape feature emerged as important, regardless of species’ dispersal syndromes: steepness. Although a steeper topography increased seed accumulation in the matrix for all dispersal syndromes, the magnitude of this effect differed among dispersal syndromes (significant three-way interaction between dispersal syndrome, steepness, and poly(distance, 2) into the matrix (*χ2* = 20.34, *P* < 0.001, Table 1)). Steep slopes increased the dispersal distance of self-dispersing seeds the most and animal-dispersed seeds the least (Table 1: ‘contrasts’).

**Table 1.** For each dispersal syndrome, we highlight two scenarios representing both steep slopes (=6°) and shallow slopes (=1°) to compare their seed abundance

| Dispersal syndrome | $X^2$ contrasts (seed abundance of steep slopes – shallow slopes) | $P$-value (comparing steepness) |
| --- | --- | --- |
| Self-dispersing | 41.22 | <0.0001* |
| Animal | 8.53 | 0.014* |
| Wind | 15.03 | 0.0005* |

*Notes:* ‘ $X^2$ contrasts’ and $P$ -values comparing steepness evaluate whether increasing steepness levels significantly increase seed abundances into the matrix (one value for each dispersal syndrome). Self-dispersing species represented 28 species in our regional community, 33 were animal-dispersed, and 14 wind-dispersed (model = abundance ~ distance\*dispersal syndrome\*steepness + (1|patch id) + (1|transect id)) on a total of 441 observations (self-dispersing = 145, animal = 233, wind = 63). For results on adult abundance change by dispersal syndrome see Appendix S2.

### Spillover of a matrix specialist resulted in either source populations or absences, not transient sink populations (hyp. iii)

We found support for the hypothesis that individuals of *Avena* spill into habitat patches close to the habitat boundary, where they also appear to persist as a source population (mean *per capita* seed production, a component of population growth, ≥1), and rarely in the centres of patches, where population growth is below replacement (mean *per capita* seed production <1, Fig. 2; significant main effect of poly(distance,2) on *Avena* abundance (*χ2* = 28.55, *P* < 0.001) and *per capita* seed production (*χ2* = 6.41, *P* = 0.041; Fig. 2B). Specifically, 1 m into patches, there were on average 98.3 seeds and adults (*range* = 0-1096, *n*_*plot*_ = 60) of *Avena* per plot, each of which produced 3.29 seeds on average (*range* = 0-9, *n*_*ind*_ =16), whereas beyond 8 m, the average abundance was 1.03 (*range* = 0-21, *n*_*plot*_ = 20), with individuals producing 1.00 seed on average (*range* = 0-2, *n*_*ind*_ = 6). Unlike patch specialists, *Avena* does not appear to accumulate a persistent seedbank in habitat patches, in contrast to a large seedbank in the matrix, suggesting that dormancy does not help persistence in unsuitable habitat. In fact, the seedbank appeared to be more sensitive to position relative to the habitat edge than did adult abundances (significant interaction between poly(distance, 2) and life stage: *χ2* = 18.70, *P* < 0.001; Fig. 2A), again, in contrast to what we observed with patch specialists (Fig. 2A vs. Fig. 1B). This significant interaction was driven by a difference in the magnitude of decline in abundance at increasing distances into patches, as both adults and seedbank declined (main effect of poly(distance, 2): *χ2* = 28.55, *P* < 0.001). Patch area (m^2^) was the only other fixed effect included in the final model but had no detectable effect on abundance (*χ2* = 6.97, *P* = 0.54) despite past work suggesting that spillover is more pronounced in small patches (Harrison et al. 2001). No other landscape factors improved the fit of either the models (all ΔAICc<2), meaning that the presence of dispersal vectors did not explain variability in *Avena* total abundance or fitness with distance into the patch.

**Figure 2.**
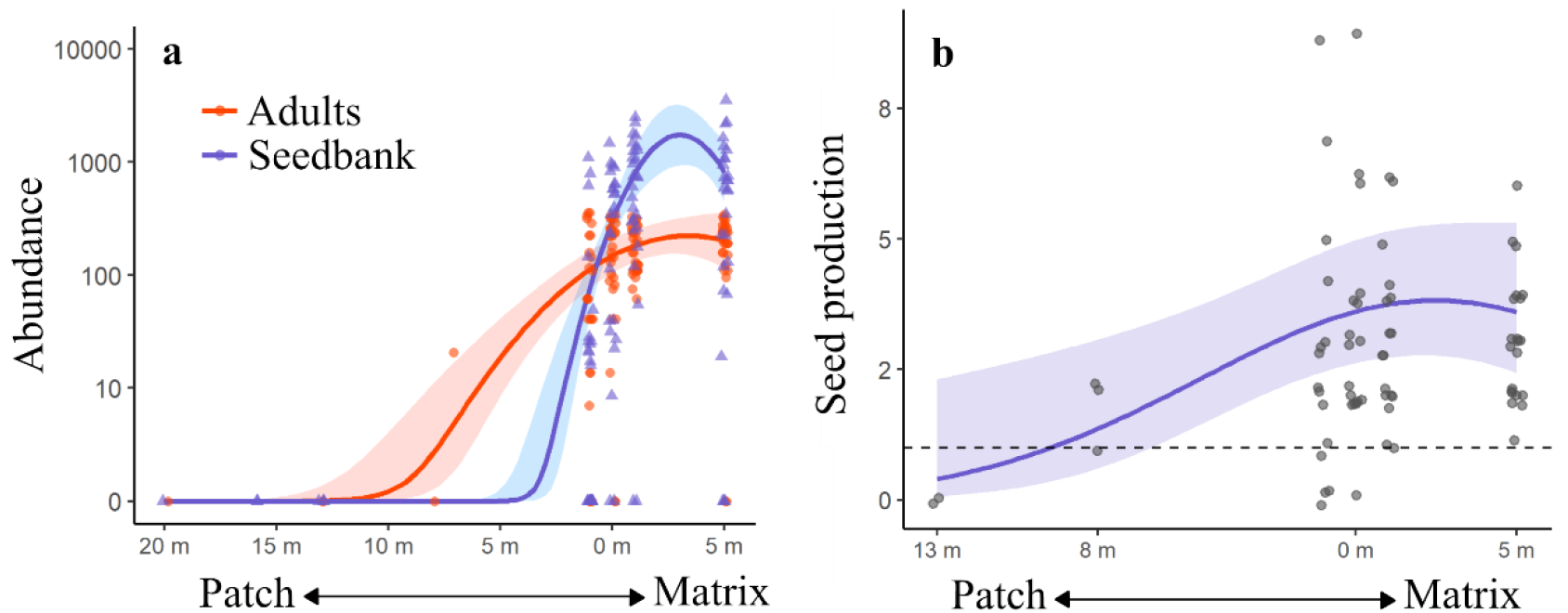
*Avena fatua*’s a) abundance and b) individual seed production declines with increasing distances into serpentine patches. a) In the matrix, there are more seeds than adults, indicating presence of a seedbank, whereas seeds and adults are similarly abundant within patches, indicating a lack of dormancy, n_obs_ = 260. We found an adult of *A. fatua* in only a single center plot (8 m into the patch). b) Individual seed production of *A. fatua* declines for individuals within patches, *n*_*ind*_ = 75. The dashed line at fitness = 1 is the point at which adults produce enough seed for replacement. Our abundance and fitness data results will have most *A. fatua* individuals in common, although some are unique to the fitness data. Points are the raw data, lines are fitted relationships from the models, and shaded areas are the 95% confidence bands of the models.

## Discussion

Dispersal is critical to biodiversity maintenance, but in patchy systems, is only achieved if individuals successfully traverse unsuitable habitat. For passively dispersing organisms, most individuals that leave a patch will fall short of reaching new ones (Bullock et al. 2017) and, in most models (e.g., (Pulliam 2000)), are assumed to die. By contrast, in our serpentine grassland system, we find that remaining dormant may be key to traversing unsuitable matrix habitat, maintaining diversity at distances far beyond species’ intrinsic dispersal abilities. We also find that the rates at which species cross habitat boundaries, and the demographic fates of those individuals when they do, do not simply mirror each other when contrasting dispersal of patch specialists in matrix habitat vs. matrix specialists in patches. Below, we first discuss our findings of patch specialists dispersing into unsuitable matrix habitat, then the dispersal and persistence of a dominant matrix species (*Avena fatua*) into patches. We then extend our findings to how they apply to patchy systems in general.

We found support for our hypothesis that dormancy helps maintain the survival of patch specialists in the matrix through one measure of diversity (i.e., total abundance) but not through the other (i.e., species richness), a mismatch which holds biological meaning (Chase et al. 2018). Let us illustrate this meaning by first reviewing for the reader what a change in species richness represents: the complete absence of every single individual of a given species. In the context of persistence across habitat boundaries, we predicted that species richness would decline most strongly for the adult community than for the seedbank (due to dormancy), meaning that for some species, every single individual that dispersed would either fail to germinate or fail to survive (i.e., species present in the seedbank but not as adults). Our findings suggest that the consequences of landing in the matrix are not so extreme as to reduce species richness (Fig. 1A), despite non-significant trends in this expected direction, but were clearly strong enough to cause a sharp decline in the total abundance of adults (Fig. 1B). Our study adds much-needed evidence for the role dormant life stages play in overcoming the challenge of dispersing through patchy landscapes (Ricketts 2001, Honnay et al. 2008, Manna et al. 2017), relevant to plants (as shown here) but also to other taxa with dormant life stages (e.g., bacteria; Wisnoski et al. 2020).

Additionally, we contend that much diversity (richness and abundance) that exists in ecological communities may lurk in the shadows, hidden as inconspicuous life forms (Pärtel 2014). Future work is needed to examine the consequences of dormancy for the permeability of unsuitable matrix habitat, as a possible means of enhancing habitat connectivity, and thus, metapopulation persistence (Casanova 2015, Wisnoski et al. 2019).

The presence of some patch specialists in the matrix as adults is consistent with past work suggesting that stepping-stone dispersal can connect populations in fragmented systems (Coughlan et al. 2017, Pedley & Dolman 2020). Although we do not have the data necessary to verify the growth rates of these adult populations, our discussion of stepping-stone dispersal applies so long as even a single individual produces any seeds. Stepping-stone dispersal through matrix habitat is the process by which a small subset of seeds that land in the matrix germinate to produce adults (typically with low fecundity), allowing newly-produced seeds to disperse a bit farther through the matrix until individuals reach new patches some number of generations later (Gilbert & Levine 2013, Van Rossum & Triest 2012). Although we do not explicitly test and compare alternative methods of connectivity (i.e., dispersal of dormant seeds vs. stepping-stone dispersal), our study provides the motivation for stepping-stone models to also consider the dispersal of dormant seeds, and in turn the possibility that these two methods act in tandem. Dispersal of dormant seeds would be especially relevant between populations when the distance between patches is greater than species’ maximum intrinsic dispersal distances, as is common in our system (Appendix S1: Table S1).

Learning whether transient populations exist in unsuitable habitats is not sufficient evidence alone to determine if transients are worth considering their contribution to metapopulation persistence. Rather, we also must understand whether transient individuals are likely to disperse through unsuitable habitat over time, eventually ending up in suitable habitat and thereby contributing to persistent population dynamics. Although dispersal distances are often described as fixed characteristics of species (Beckman et al. 2018), we found that those distances were stretched in some patches and not others depending on landscape topography (Dunning et al. 1992, Lowe & McPeek 2014). Specifically, we found that seeds accumulated most in the matrix when patches were on a steep slope, compared to patches that were on a shallow slope (Table 1). Our results suggest that dormant seeds with different primary dispersal adaptations (e.g., wind, animals) can secondarily move with material flows in our grassland system, suggesting that physical landscape features may be a strong driver of seed dispersal through the matrix. Thus, seeds may experience similarities in how landscape connectivity is manifested. Therefore, upslope habitat, particularly when biodiverse, may act as source patches for many species dispersing into downslope patches (Mouquet et al. 2013), similar to processes in dendritic riverine systems (Carrara et al. 2012, Little & Altermatt 2018).

In contrast to the findings of our study, previous research in the same system has shown that dispersal vectors do influence how species with different dispersal syndromes experience connectivity, using parameterizations of connectivity models (Germain et al. 2019). This disconnect between previous and current research might arise as a consequence of differences in the spatial scale of inference implicit in each study: seedbank assays taking place within 5 m of patches (this study) vs. statistical estimates of connectivity at all distances between patches (range = 0.75-356 m; (Germain et al. 2019)). The shape of the dispersal kernel, and thus degrees of habitat connectivity, might be shaped by different mechanisms at two scales (Rogers et al. 2019): by slope on small scales for all species, shifting the bulk density (i.e., frequent, short-distance dispersal) of the kernel regardless of dispersal syndrome, and by dispersal vectors on large scales, stretching the kernel’s tail (i.e., infrequent, long-distance dispersal).

By contrasting our findings between patch specialists in the matrix and a matrix specialist (*Avena fatua*) in patches, it is clear that the demographic fates, and thus, ecological consequences, of individuals crossing the habitat boundary are not symmetric in both directions. Given that *Avena* was so abundant in matrix habitat, we expected that it would form abundant sink populations in serpentine patches, such like patch specialists in the matrix—we were wrong in two ways. First, up to a few metres into patches, *Avena* did form abundant populations (∼100 individuals per quadrat), but these populations were source populations (mean seed production ≥ 1), not sink populations (mean seed production < 1). We were surprised that patch edges were as suitable to *Avena* as they were. We suggest that this surprise is the result of the presence of fuzzy boundaries between serpentine patches and the non-serpentine matrix, where the negative effects of an abiotically-harsh serpentine soil environment are counteracted by positive effects of weakened intraspecific competition (Germain et al. 2018). This would explain the difference between the number of seeds and the number of adults differed we observed in the matrix, compared to in patches (Fig. 2A): the germination and removal of seedlings due to competition (Goldberg et al. 2001).

Second, deeper into patches, such as near patch centres, *Avena* individuals were largely absent. The few individuals that were found near the patch centres hovered at population replacement, which, because there are so few individuals, means that stochastic extinction is probable (Gravel et al. 2011). In other words, abundances of *Avena* were concentrated in suitable habitat and not in unsuitable habitat (Shoemaker & Melbourne 2016), suggesting that, unlike for patch specialists, dispersal and dormancy (if any; Naylor & Jana 1976) were not strong enough to maintain a biologically-relevant population of *Avena*. Why would *Avena* not evolve strong dormancy and dispersal given the obvious negative impacts of patch conditions on reproductive output (i.e., 4x lower than in the matrix)? We contend that the answer is that crossing unsuitable habitat is less crucial for the success of matrix specialists, who are instead free to permeate through continuous habitat unencumbered. Overall, our findings align with past research suggesting that spillover of invasive species will be most detrimental in small patches, due to high abundances at the habitat boundary (Harrison et al. 2001). However, given how few seeds and adults were found more deeply within patches, serpentine patches do appear to act as effective habitat refuges from invaders.

## Conclusions

We set out to elucidate the demographic fate of dispersers that fall short of suitable habitat, specifically, whether dormancy and sink populations allow the transient persistence of species across habitat boundaries. Dispersal through inhospitable conditions within patchy systems is critical for habitat connectivity, which in turn maintains both local and regional biodiversity (Freestone & Inouye 2006, Germain et al. 2017). In a fragmented landscape of serpentine patches separated by a continuous, invaded matrix habitat, we found that dormancy is a process that allows native biodiversity to commonly survive in unsuitable matrix habitat. Invasive matrix species, on the other hand, rarely persist in unsuitable patch habitat, and are found as small sink populations when they do. Our work informs biodiversity science by motivating the inclusion of the movement of dormant life stages in future connectivity models, towards better predictions of biodiversity change in fragmented landscapes.

## Acknowledgments

We thank Kately Nikiforuk, Jenny Mackay, Catherine Koehler, Mia Waters, and Dennis Chiu for their assistance with field work and data collection. Special thanks to Susan Harrison for lending us her long-term data. Thanks to Chelsea Little and Jesse Fleri for assisting with spatial analyses and helpful suggestions on the manuscript. This work would not have been possible without financial support from NSERC Discovery Grant (2019-04872), Dept. Zoology.

## Author contributions

MCS and RMG both conceived the ideas, designed methodology, and collected the data. MCS analysed the data and led the writing of the manuscript. RMG secured funding for the project and provided guidance throughout the analysis and writing.

