## Appendices S1 & S2 for "Dispersal across habitat boundaries: uncovering the demographic fates of populations in unsuitable habitat"

APPENDIX S1

*Additional species information*

**Table S1.** The scientific name of all identified species of our study. Dispersal mode was obtained from Jones et al. (2015). Species status where obtained from CABI.org. Habitat were determined using a historical dataset spanning 17 years within the greater reserve area of our specific study (S. Harrison). Sensitivity level refers to the strength of species association to their primary habitat, where 1 = very associated due to much high abundance in one habitat, to 3 = loosely associated to due to rarity in both habitats. our analysis uses patch specialists in sensitivity level 1 and 2.

| Family | Species name | Status | Primary dispersal mode | Habitat affinity | Sensitivity level |
|---|---|---|---|---|---|
| *Asteraceae* | *Achyrachaena mollis* | native | Animal | patch | 1 |
| *Fabaceae* | *Acmispon brachycarpus* | native | self-dispersing | patch | 1 |
| *Fabaceae* | *Acmispon wrangelianus* | native | self-dispersing | patch | 1 |
| *Asteraceae* | *Agoseris heterophylla* | native | Wind | patch | 1 |
| *Amaryllidaceae* | *Allium amplectens* | native | Self-dispersing | patch | 1 |
| *Amaryllidaceae* | *Allium fimbriatum* | native | Self-dispersing | patch | 1 |
| *Primulaceae* | *Anagallis arvensis* | naturalized | Animal | patch | 2 |

| | | | | | |
|---|---|---|---|---|---|
| *Asteraceae* | *Ancistrocarphus filagineus* | native | Animal | patch | 1 |
| *Brassicaceae* | *Athysanus pusillus* | native | Animal | matrix | |
| *Poacaea* | *Avena sp.* | invasive | Animal | matrix | |
| *Poacaea* | *Bromus hordeaceus* | invasive | Animal | matrix | |
| *Poacaea* | *Bromus madritensis* | invasive | Animal | patch | 1 |
| *Montiaceae* | *Calandrinia ciliata* | native | Self-dispersing | patch | 2 |
| *Liliaceae* | *Calochortus luteus* | native | Wind | patch | 1 |
| *Asteraceae* | *Calycadenia pauciflora* | endemic | Wind | patch | 1 |
| *Onagraceae* | *Camissonia graciliflora* | native | Self-dispersing | patch | 1 |
| *Brassicaceae* | *Cardamine oligosperma* | native | Self-dispersing | rare | |
| *Orobanchaceae* | *Castilleja attenuata* | native | Wind | patch | 1 |
| *Asteraceae* | *Centaurea solstitialis* | invasive | Wind | matrix | |
| *Onagraceae* | *Clarkia gracilis* | native | Self-dispersing | patch | 1 |
| *Plantaginaceae* | *Collinsia sparsiflora* | native | Self-dispersing | patch | 1 |
| *Asteraceae* | *Croton setiger* | native | Wind | patch | 1 |

| | | | | | |
|---|---|---|---|---|---|
| *Asparagaceae* | *Dichelostemma capitatum* | native | Animal | patch | 1 |
| *Poaceae* | *Elymus caput-medusae* | invasive | Animal | matrix | |
| *Polygonaceae* | *Eriogonum nudum* | native | Animal | patch | 1 |
| *Asteraceae* | *Eriophyllum lanatum* | native | Self-dispersing | patch | 3 |
| *Geraniaceae* | *Erodium cicutarium* | invasive | Animal | matrix | |
| *Euphorbiaceae* | *Euphorbia crenulata* | native | Self-dispersing | patch | 3 |
| *Poaceae* | *Festuca myuros* | invasive | Animal | matrix | |
| *Rubiaceae* | *Galium aparine* | native | Animal | matrix | |
| *Poaceae* | *Gastridium ventricosum* | naturalized | Animal | rare | |
| *Polemoniaceae* | *Gilia capitata* | native | Self-dispersing | patch | 3 |
| *Campanulaceae* | *Githopsis specularioides* | native | Self-dispersing | rare | |
| *Asteraceae* | *Grindelia camporum* | endemic | Self-dispersing | patch | 1 |
| *Asteraceae* | *Hemizonia congesta* | native | Self-dispersing | patch | 1 |
| *Linaceae* | *Hesperolinon sp.* | endemic | Self-dispersing | patch | 1 |
| *Poaceae* | *Hordeum sp.* | invasive | Animal | patch | 3 |

| | | | | | |
|---|---|---|---|---|---|
| *Asteraceae* | *Hypochaeris glabra* | invasive | Wind | matrix | |
| *Asteraceae* | *Lactuca sp.* | naturalized | Wind | patch | 2 |
| *Asteraceae* | *Lasthenia californica* | native | Animal | patch | 1 |
| *Brassicaceae* | *Lepidia nitidum* | native | Self-dispersing | patch | 2 |
| *Polemoniaceae* | *Linanthus bicolor* | native | Self-dispersing | patch | 2 |
| *Asteraceae* | *Logfia californica* | native | Animal | patch | 1 |
| *Asteraceae* | *Logfia gallica* | naturalized | Animal | matrix | |
| *Poaceae* | *Lolium multiflorum* | naturalized | Animal | patch | 3 |
| *Fabaceae* | *Lupinus succulentus* | native | Self-dispersing | patch | 1 |
| *Asteraceae* | *Madia exigua* | native | Wind & Animal | matrix | |
| *Fabaceae* | *Medicago polymorpha* | invasive | Animal | matrix | |
| *Fabaceae* | *Melilotus indica* | naturalized | Animal | rare | |
| *Asteraceae* | *Micropus californicus* | native | Self-dispersing | patch | 1 |
| *Asteraceae* | *Microseris douglasii* | native | Wind | patch | 1 |
| *Phrymaceae* | *Mimulus douglasii* | native | Self-dispersing | patch | 1 |

| | | | | | |
|---|---|---|---|---|---|
| *Phrymaceae* | *Mimulus nudatus* | native | Animal | patch | 3 |
| *Caryophyllaceae* | *Minuartia douglasii* | native | Self-dispersing | patch | 1 |
| *Poacaea* | *Nassella pulchra* | native | Animal | patch | 1 |
| *Polemoniaceae* | *Navarretia jepsonii* | endemic | Animal | patch | 1 |
| *Polemoniaceae* | *Navarretia pubescens* | native | Animal | matrix | |
| *Caryophyllaceae* | *Petrorhagia prolifera* | naturalized | Wind | matrix | |
| *Polemoniaceae* | *Phlox gracilis* | native | Wind | patch | 2 |
| *Plantaginaceae* | *Plantego erecta* | native | Animal | patch | 1 |
| *Poacaea* | *Poa secunda* | native | Self-dispersing | patch | 1 |
| *Asteraceae* | *Rigiopappus leptocladus* | native | Wind | patch | 1 |
| *Malvales* | *Sidalcea diploscypha* | endemic | Animal | patch | 1 |
| *Asteraceae* | *Sonchus oleraceus* | naturalized | Wind | insufficient evidence | |
| *Caryophyllaceae* | *Stellaria nitens* | native | Self-dispersing | patch | 1 |
| *Fabaceae* | *Trifolium albopurpureum* | native | Self-dispersing | patch | 1 |
| *Fabaceae* | *Trifolium bifidum* | native | Self-dispersing | matrix | |

| | | | | | |
|---|---|---|---|---|---|
| *Fabaceae* | *Trifolium fucatum* | native | Self-dispersing | patch | 1 |
| *Fabaceae* | *Trifolium microdon* | endemic | Animal | matrix | |
| *Asparagaceae* | *Triteleia laxa* | endemic | Self-dispersing | matrix | |
| *Asteraceae* | *Uropappus lindleyi* | native | Wind | patch | 3 |
| *Fabaceae* | *Vicia villosa* | invasive | Self-dispersing | matrix | |
| *Poaceae* | *Vulpia bromoides* | naturalized | Wind & Animal | rare | |
| *Poaceae* | *Vulpia microstachys* | native | Animal | patch | 1 |
| *Liliaceae* | *Zigadenus fremontii* | native | Self-dispersing | patch | 1 |

*Sensitivity analysis*

To determine what species were associated with serpentine patches, we conducted a sensitivity analysis using three levels of association. If species were either not clearly associated with any single habitat or were rare in both, we labelled them as “unassigned” and did not include them in a specialist community. The most conservative assemblage for patch specialists included only those whose abundance differed by >100 individuals between habitat types. The mid-conservative assemblage contained the same species, but included those that were rare in both habitats, but proportionally were >10% more abundant in the serpentine site ranks. The least-conservative species included all the above species, but included species that ranked rare (0-50 individuals), and were found relatively more frequently in serpentine sites (Appendix S1: Table S1, S2).

**Table S2.** Patch sensitivity model output. The three levels of sensitivity are 1 being most conservative, containing the least amount of species, and 3 being least-conservative, including the most species. For build of the selected models refer to Appendix S2.

| **Sensitivity level** | **Analysis** | **Selected model** | **$X^2$** | ***P*-value** | ***N*-species** |
|---|---|---|---|---|---|
| 1 | Richness | A4 | 0.54 | 0.7623 | 38 |
| 2 | Richness | A4 | 0.44 | 0.8007 | 44 |
| 3 | Richness | A4 | 0.46 | 0.7923 | 51 |
| 1 | Total abundance | B2 | 7.95 | 0.01877* | 38 |
| 2 | Total abundance | B2 | 9.02 | 0.01101* | 44 |
| 3 | Total abundance | B2 | 8.62 | 0.01346* | 51 |

APPENDIX S2

*Model selection*

**Failed dispersal into the matrix results in a transient dormant seedbank (hyp. i)**

**Table S1**. A) Stepwise model selection for richness, and B) for patch specialist total abundance. Bolded model indicates the final model with the lowest AICc.

| Explanatory ~ Predictor | Distribution | Model | New terms | AICc | ΔAICc |
|---|---|---|---|---|---|
| | A. RICHNESS | | | | |
| richness ~ exp(distance)* life-stage | poisson | 1 | -- | 1006.35 | 0 |
| **richness ~ poly(exp(distance),2)* life-stage** | **poisson** | **2** | **--** | **990.20** | **16.15** |
| | TOTAL ABUNDANCE | | | | |
| total abundance ~ exp(distance)*life-stage | beta | 1 | -- | -232.43 | 0 |
| **total abundance ~ poly(exp(distance),2)* life-stage** | **beta** | **2** | **--** | **-237.29** | 4.86 |

Notes: Each base model included the interaction between distance over habitat boundaries and life-stage (germination trial and in-field surveys) as it was central to our biological question of whether the slope of the seedbank differed from the slope of adults for different community metrics. Each model contained the random effects (1|patch) and (1|transect) to control for our nested sampling design (not shown in table). For the richness variable, the predictor distance is exponentiated to account for underfitting of low values. The measure of total abundance we use for B includes the summed abundance of each species found within plots. As such it represents the number of patch individuals found across habitats. Abundance is relative within life stages because it was scaled so that the recorded abundance of the germination trial and the in-field adult survey could be interpreted on the same scale, even though the survey area differed between those life-stages. For models using patch specialists only (richness and total abundance), we included patch species that had a sensitivity level of 1 & 2 (mid-conservative measure, see Appendix S1: Table S1).

*Richness and total abundance predicted by categorical positions*:

I performed the same analyses as reported above for patch specialist richness and total abundance, but to test whether the continuous 'distance from boundary' models constrained the shape of life stage slopes, here we use distance as a categorical variable 'position'. Species richness of patch specialists declined by increasing position into the matrix (main effect: $\chi^2$ = 38.36, $P < 0.001$) a pattern that did not differ between life stages (position x lifestage: $\chi^2$ = 2.05, $P = 0.72$). Total abundance of patch specialists differed between positions (main effect: $\chi^2$ = 35.70, $P < 0.001$), a pattern that differed between life stages (position x lifestage: $\chi^2$ = 12.93, $P$ = 0.012). These results using distance as a categorical variable 'position' quantitatively mirror the patterns reported in the main text that used distance as a continuous variable.

**Table S2.** A) Stepwise model selection for richness, B) for patch specialist abundance when distance is categorized by position. Bolded model indicates the final model with the lowest AICc.

| Explanatory ~ Predictor | Distribution | Model | New terms | AICc | ΔAICc |
|---|---|---|---|---|---|
| A. | RICHNESS | | | | |

| | | | | | |
|---|---|---|---|---|---|
| richness ~ position* life-stage | poisson | 1 | -- | 997.58 | 0 |
| richness ~ position* life-stage | poisson | 2 | transect type | 1000.01 | -2.43 |
| **richness ~ position* life-stage** | **poisson** | **3** | **steepness** | **961.58** | 36 |
| richness ~ position* life-stage | poisson | 4 | steepness* life-stage | 963.78 | -2.2 |
| richness ~ position* life-stage | poisson | 5 | steepness* position | 968.49 | -6.88 |
| | | B. TOTAL ABUNDANCE | | | |
| **Total abundance ~ position*life-stage** | **beta** | **1** | **--** | **-232.80** | 0 |
| Total abundance ~ position*life-stage | beta | 2 | transect type | -230.74 | -2.06 |
| Total abundance ~ position*life-stage | beta | 3 | steepness | -216.11 | 16.69 |

Notes: Each base model included the interaction between distance over habitat boundaries and life-stage (germination trial and in-field surveys) as it was central to our biological question of whether the seedbank differed from the adults in different positions for richness and total abundance. Each model contained the random effects (1|patch) and (1|transect) to control for our nested sampling design (not shown in table). The measure of total abundance we use for B includes the summed abundance of each species found within plots. As such it represents the number of patch individuals found across local habitat. Abundance is relative because it was scaled so that the recorded abundance of the germination trial and the in-field adult survey could be interpreted on the same scale, even though the survey area differed between those life-stages. For models using patch specialists only (richness and total abundance), we included patch species that had a sensitivity level of 1 & 2 (mid-conservative measure, see Table S2).

*Seed dispersal is extended by interacting syndromes and vectors (hyp. ii)*

**Table S3.** AICc Stepwise model selection for seed dispersal syndrome models. Bolded model indicates the final model with the lowest AICc.

| Explanatory ~ Predictor | Distribution | Model | New terms | AICc | ΔAICc |
|---|---|---|---|---|---|
| Seed abundance ~ distance | poisson | 1 | -- | 8115.99 | 0 |
| Seed abundance ~ poly(distance,2) | poisson | 2 | -- | 8095.38 | 20.61 |
| Update .~. 2 | poisson | 3 | dispersal mode | 7790.73 | 304.65 |
| Update .~. 2 | poisson | 4 | transect type | 8100.00 | -4.62 |
| Update .~. 2 | poisson | 5 | steepness | 8013.21 | 82.17 |
| Update .~. 3 | poisson | 6 | dispersal mode + steepness | 7693.74 | 96.99 |
| Update .~. 6 | poisson | 7 | poly(distance,2)* dispersal mode | 7616.26 | 77.48 |
| Update .~. 6 | poisson | 8 | dispersal model*steepness | 7634.31 | 59.43 |
| Update .~. 7 | poisson | 9 | dispersal mode*steepness | 7609.84 | 6.42 |
| **Update .~. 9** | **poisson** | **10** | **poly(distance,2)* dispersal mode* steepness** | **7598.15** | 11.69 |

Notes: Each model contained the random effects (1|patch) and (1|transect) to control for our nested sampling design (not shown in table).

*Matrix species spillover results in transient sink populations (hyp. iii)*

**Table S4**. The model selection methods for *Avena fatua* A) abundance and B) seed production across habitat boundaries. Bolded model indicates the final model with the lowest AICc.

| **Explanatory ~ Predictor** | **Distribution** | **Model** | **New terms** | **AICc** | **ΔAICc** |
|---|---|---|---|---|---|
| | A. ABUNDANCE | | | | |
| Abundance ~ distance* life-stage | neg. binom | 1 | -- | 3021.07 | 0 |
| Abundance ~ poly(distance,2)* life-stage | neg. binom | 2 | -- | 2980.55 | 40.52 |
| **Update .~. 2** | **neg. binom** | **3** | **area** | **2798.46** | 182.09 |
| Update .~. 2 | neg. binom | 4 | transect type | 2980.45 | 0.1 |
| Update .~. 2 | neg. binom | 5 | steepness | Singular fit | |
| Update .~. 2 | neg. binom | 6 | ndvi | Singular fit | |
| Update .~. 3 | neg. binom | 7 | area*life-stage | 2798.89 | 0.4 |
| Update .~. 3 | neg. binom | 8 | area*poly(distance,2) | 2800.88 | -2.42 |
| | 2. INDIVIDUAL SEED PRODUCTION | | | | |
| Seeds produced ~ distance | poisson | 1 | -- | 298.10 | 0 |

| | | | | | |
|---|---|---|---|---|---|
| **Seeds produced ~ poly(distance,2)** | **poisson** | **2** | **--** | **294.46** | 3.64 |
| Update .~. 2 | poisson | 3 | transect type | 298.84 | -4.38 |
| Update .~. 2 | poisson | 4 | steepness | 295.25 | -0.79 |

Notes: Each model contained the random effects (1|patch) and (1|transect) to control for our nested sampling design (not shown in table).

**Table S5.** Here we compare full models of each response variable to the selected final model as a check as recommended by Mundry and Nunn (2009). Final models were selected for with forward stepwise procedure (Appendix S2: Table S1, S3, S4). Chi-squared results are given from the Anova type III command from the 'car' package. *P*-values are reported only for the interaction term. The full and final models are qualitatively similar for all response variables.

| **Model type** | **Response ~ Predictors** | $X^2$ | ***P*-value** |
|---|---|---|---|
| Final | Richness ~ poly(exp(distance),2)* life-stage | 0.44 | 0.801 |
| Full | Richness ~ poly(exp(distance),2)* life-stage | 0.41 | 0.816 |
| Final | Total abundance ~ poly(exp(distance),2)* life-stage | 9.03 | 0.011 |
| Full | Total abundance ~poly(exp(distance),2)* life-stage | 9.69 | 0.001 |

| | | | |
|---|---|---|---|
| Final | Seed abundance ~ poly(distance,2)* dispersal syndrome* steepness | 20.39 | <0.001 |
| Full (#1) | Seed abundance ~ poly(distance,2)* dispersal syndrome * steepness + transect type | 20.47 | <0.001 |
| Full (#2) | Seed abundance ~ poly(distance,2)* dispersal syndrome * transect type + steepness | 112.22 | <0.001 |
| Final | Avena abundance ~ poly(distance,2)* life-stage + area | 18.70 | <0.001 |
| Full | Avena abundance ~ poly(distance,2)* life-stage + steepness + transect type + area + ndvi | Non-convergent | --- |
| Final | Avena fitness ~ poly(distance,2) | 6.41 | 0.041 |
| Full | Avena fitness ~ poly(distance,2) + transect type + steepness | 6.78 | 0.034 |

Notes: All models contain the random effects 1|transect id and 1|patch id. Seed abundance has two full models as it is not biologically feasible for transect type and steepness to interact with one another.
